# Matrix stiffness and stress relaxation regulate osteogenesis through histone demethylases KDM4B and KDM6B

**DOI:** 10.1101/2025.07.16.665229

**Authors:** Ian M. Tayler, Amy Zhu, Ryan S. Stowers

**Affiliations:** Dept. of Molecular, Cellular and Developmental Biology, UC Santa Barbara; Dept. of Mechanical Engineering, UC Santa Barbara; Dept. of Bioengineering, UC Santa Barbara

## Abstract

Stem cells sense biophysical cues within their extracellular microenvironment and respond via mechanotransduction signaling pathways that induce changes in gene expression and associated cell fate outcomes. Histone modifying enzymes are known to drive stem cell differentiation through changes in chromatin accessibility, but little is understood as to how extracellular matrix (ECM) mechanics regulate epigenomic remodeling. Here, we utilize alginate hydrogels with tunable mechanical properties to investigate the role of both matrix stiffness and viscoelasticity on histone demethylase expression and activity during osteogenic differentiation of human bone marrow-derived mesenchymal stem cells (hBMSCs). Our results reveal that the expression of two histone demethylases, KDM4B and KDM6B, are upregulated during osteogenesis in response to stiff, viscoelastic matrix conditions. Inhibition of mechanotransduction signaling pathways reduces expression of KDM4B and KDM6B and hinders osteogenic differentiation overall. Interestingly, phosphorylation of SMAD 1/5/8 was shown to increase in cells cultured in stiff, stress relaxing matrices, and pharmacological inhibition of SMAD 1/5/8 activation reduced expression of KDM4B and KDM6B and decreased osteogenic differentiation. Taken together, our results reveal novel impacts of stem cell mechanotransduction signaling events that promote osteogenesis through epigenetic remodeling.

## Introduction

Bone marrow-derived mesenchymal stem cells (BMSCs) are multipotent, mechanosensitive progenitors capable of differentiating into numerous cell types, including chondrocytes, adipocytes, and osteoblasts. During bone remodeling and fracture repair, BMSCs undergo osteogenic differentiation in response to sensing microenvironmental mechanical properties (Darnell *et al*., 2017; Duan and Lu, 2021; Woloszyk *et al*., 2022). In vitro experiments have shown that extracellular matrix (ECM) mechanical properties such as stiffness and stress relaxation regulate gene expression (Hwang *et al*., 2015; Darnell *et al*., 2018) and ultimately stem cell differentiation through activation of mechanotransduction pathways (Engler *et al*., 2006; Huebsch *et al*., 2010; Parekh *et al*., 2011; Chaudhuri *et al*., 2016; Lee *et al*., 2019). During osteogenesis, mechanosignaling emanates from integrin-based adhesion complexes and mechanically-sensitive ion channels to induce cytoskeletal remodeling, drive actomyosin contractility, and activate downstream signaling pathways such as RhoA, PI3K, and YAP/TAZ. (Fujita *et al*., 2004; McBeath *et al*., 2004; Taubenberger *et al*., 2010; Huang *et al*., 2023; Liu *et al*., 2023; Li *et al*.). Intriguingly, it has become increasingly evident that ECM mechanics can induce epigenetic remodeling via cytoskeletal mechanotransduction to alter cellular phenotypes (Downing *et al*., 2013; Stowers *et al*., 2019; Jang *et al*., 2021; Wu *et al*., 2025). However, it remains unclear whether mechanically induced osteogenesis is influenced by epigenetic remodeling.

Osteogenesis is regulated through canonical bone morphogen protein (BMP) signaling via BMP receptor phosphorylation of SMAD 1/5/8 transcription factors, which translocate to the nucleus and facilitate osteogenic gene expression (Rahman *et al*., 2015; Wu *et al*., 2016). In concert with canonical BMP-SMAD signaling, BMSC lineage specification occurs through associated changes in gene expression that are ultimately regulated by chromatin accessibility (Huang *et al*., 2015; Kanazawa *et al*., 2021; Walewska *et al*., 2023; Zhu *et al*., 2024). These changes in accessibility are often governed by histone-modifying enzymes that chemically modify the N-terminal tails of histone subunits, commonly by acetylation or methylation, to enhance or repress gene transcription. Previous work has demonstrated that osteogenesis is epigenetically regulated through two lysine-specific histone demethylases, KDM4B and KDM6B, that are essential for osteogenic differentiation (Ye *et al*., 2012; Xu *et al*., 2013; Deng *et al*., 2022; Liu *et al*., 2022). Expression of KDM4B and KDM6B is upregulated in BMSCs upon treatment with BMP 4/7 (Ye *et al*., 2012), and the loss of their expression significantly decreases osteogenic differentiation. The demethylases KDM4B and KDM6B function at H3K9me3 and H3K27me3 sites, respectively, to coordinate the de-repression of key osteogenic genes (Ye *et al*., 2012; Xu *et al*., 2013; Deng *et al*., 2022). However, it is not well understood how matrix mechanics regulate epigenetic state transitions in BMSCs during osteogenic differentiation, or whether KDM4B or KDM6B are mechanoresponsive during this lineage commitment.

In the present study, we aimed to determine how matrix mechanics impact the expression and activity of KDM4B and KDM6B in the context of mechanically induced osteogenic differentiation of human BMSCs (hBMSCs). We utilized a mechanically tunable, three-dimensional (3D) alginate hydrogel cell culture platform with independently tunable stiffness and stress relaxation rates. We have identified that stiffness and stress relaxation both regulate the expression of KDM4B and KDM6B and downstream hallmarks of early osteogenic differentiation. By encapsulating hBMSCs in these hydrogel networks, we delineated the combined effects of stiffness and stress relaxation to investigate how mechanotransduction signaling impacts epigenetic changes during osteogenesis. Here, we describe a novel function for the histone modifiers KDM4B and KDM6B and demonstrate that they are dependent on both actin cytoskeleton contractility and SMAD 1/5/8 signaling in 3D matrices. Our work provides further insight into how the cellular microenvironment impacts the lineage specification of mechanosensitive stem cells.

## Results

### Hydrogel stiffness and stress relaxation rate regulate osteogenesis in 3D

To determine the influence of ECM mechanics on epigenetic changes during early osteogenic differentiation, we utilized an alginate hydrogel system that enables independent tuning of both matrix elastic modulus and stress relaxation rate (Figure 1A). Hydrogel crosslinking densities were tuned by varying the concentration of calcium to achieve elastic moduli of 20 kPa (stiff) or 3 kPa (soft) (Figure 1B). These elastic moduli were selected based on previous reports identifying stiffness-mediated osteogenic differentiation of MSCs in 3D (Huebsch *et al*., 2010; Chaudhuri *et al*., 2016). The hydrogel stress relaxation rate was tuned independently of stiffness by using alginates of varied molecular weights (Figure 1C-D), and by conjugation of polyethylene glycol (PEG) to low molecular weight alginate polymers via carbodiimide chemistry (Nam *et al*., 2019). Additionally, the RGD adhesion motif was then coupled to all alginate polymers as previously described (Rowley *et al*., 1999) at a final concentration of 1,500 μM.

**Figure 1:**
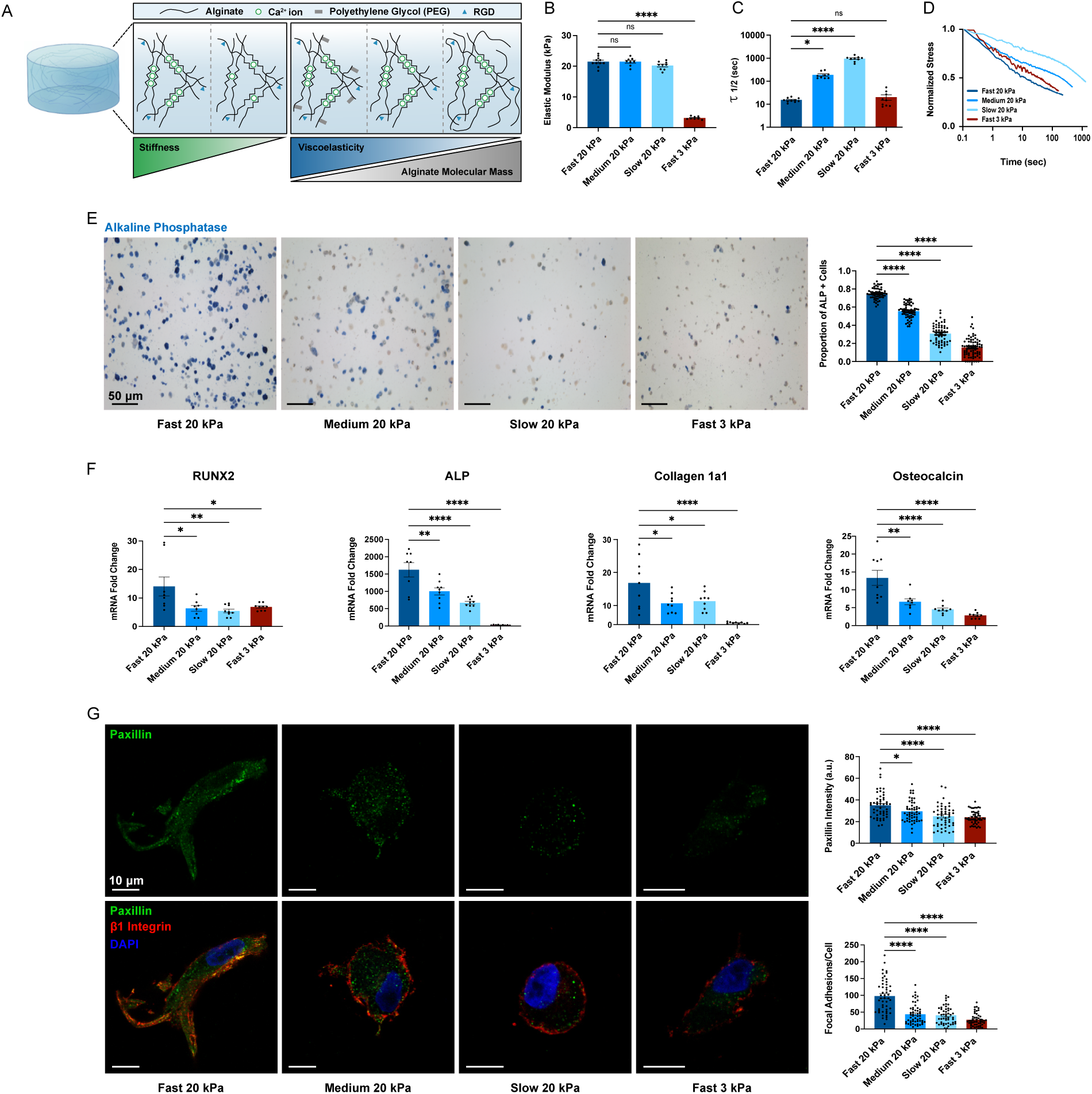
Matrix stiffness and stress relaxation direct osteogenesis in 3D. **A**, Schematic illustrating the approach to tuning alginate hydrogel mechanical properties. **B**, Measurements of hydrogel elastic modulus one day after crosslinking. **C**,**D**, Quantifications of stress relaxation half-times across hydrogel matrices. **E**, Representative images of alkaline phosphatase staining (left) indicating osteogenic differentiation of hBMSCs. Quantification of ALP-positive cells across hydrogel conditions (right). **F**, Gene expression data for early osteogenic markers from cells cultured across hydrogel conditions. **G**, Representative micrographs of cells cultured across hydrogel conditions stained for paxillin, β1 integrin, and nuclei (left) with quantifications of both paxillin intensity and colocalization of paxillin with β1 integrin (right). Statistical significance was determined by one-way analysis of variance (ANOVA) followed by Dunnett’s multiple testing correction. n = 3 replicates per condition. * indicates p < 0.05, ** p < 0.01, *** p < 0.001, **** p < 0.0001.

After characterizing the mechanical parameters of our hydrogel-based cell culture system, hBMSCs were encapsulated in alginate matrices and cultured for 7 days in osteogenic differentiation media. In line with previous studies (Chaudhuri *et al*., 2016; Lee *et al*., 2019), the cells in stiff, fast-relaxing matrices exhibited significantly more osteogenesis assessed by the proportion of cells that stained positive for alkaline phosphatase (ALP), compared to softer or slower-relaxing hydrogel conditions (Figure 1E). Further analysis of early osteogenic differentiation markers by quantitative polymerase chain reaction (qPCR) revealed that cells cultured in stiff, fast-relaxing matrices displayed significantly greater gene expression of RUNX2, ALP, Collagen 1a1, and Osteocalcin compared to all other mechanical conditions (Figure 1F). We next sought to identify the ECM-mediated signaling pathways contributing to the hBMSCs’ differential response to matrix stiffness and stress relaxation cues, both which are known to promote osteogenic differentiation (Chaudhuri *et al*., 2016; Darnell *et al*., 2017; Lee *et al*., 2019). In particular, integrin signaling and focal adhesion formation are enhanced in fast stress relaxing matrices (Lou *et al*., 2018). Furthermore, evidence from the field has demonstrated the necessity of integrin-based signaling to promote mechanically induced osteogenic differentiation (Comisar *et al*., 2007; Chen and Jacobs, 2013; Chaudhuri *et al*., 2016; Görlitz *et al*., 2024; Li *et al*., 2024a; Wan *et al*., 2024). In line with these observations, we assessed the extent of focal adhesion formation across our hydrogel conditions by quantifying the colocalization of β1 integrin and paxillin immunofluorescence signals, as well as total intensity of paxillin fluorescence. Interestingly, cells in stiff, fast-relaxing matrices had significantly more regions of colocalized signals per cell, compared to cells in stiff, slower-relaxing or soft, fast-relaxing conditions (Figure 1G). Furthermore, increased focal adhesion marker colocalization coincided with significantly greater cell spreading and decreased cell circularity in the stiff, fast-relaxing matrices, which was evaluated by changes in cell area (Supplemental Figure 1). Taken together, these results indicate that stiff, fast-relaxing matrices promote hBMSC osteogenesis through focal adhesion-mediated cell-matrix interactions.

### Stiff, fast-relaxing matrices promote increased expression of KDM4B and KDM6B

We next sought to investigate how the histone modifiers KDM4B and KDM6B are affected by matrix mechanics. Intriguingly, we found that gene expression of both KDM4B and KDM6B was significantly higher in cells cultured within stiff, fast-relaxing matrices relative to the soft or slow-relaxing hydrogel conditions (Figure 2A-B). We next aimed to characterize subcellular localization of KDM4B and KDM6B in different matrix mechanical conditions during osteogenesis by immunofluorescence staining and confocal microscopy. We found that both KDM4B and KDM6B display significantly greater nuclear fluorescence intensity in stiff, fast-relaxing matrices compared to all other matrix conditions. Additionally, we found that stiff and fast-relaxing matrices promoted the greatest ratio of nuclear-to-cytoplasmic KDM4B and KDM6B (Figure 2C-D). These results suggest that KDM4B and KDM6B accumulate within the nucleus in stiff, fast-relaxing matrix conditions to further promote osteogenic gene expression relative to soft or slow-relaxing matrices. Given that KDM4B and KDM6B are histone demethylases that act on trimethylated H3K9 and

**Figure 2:**
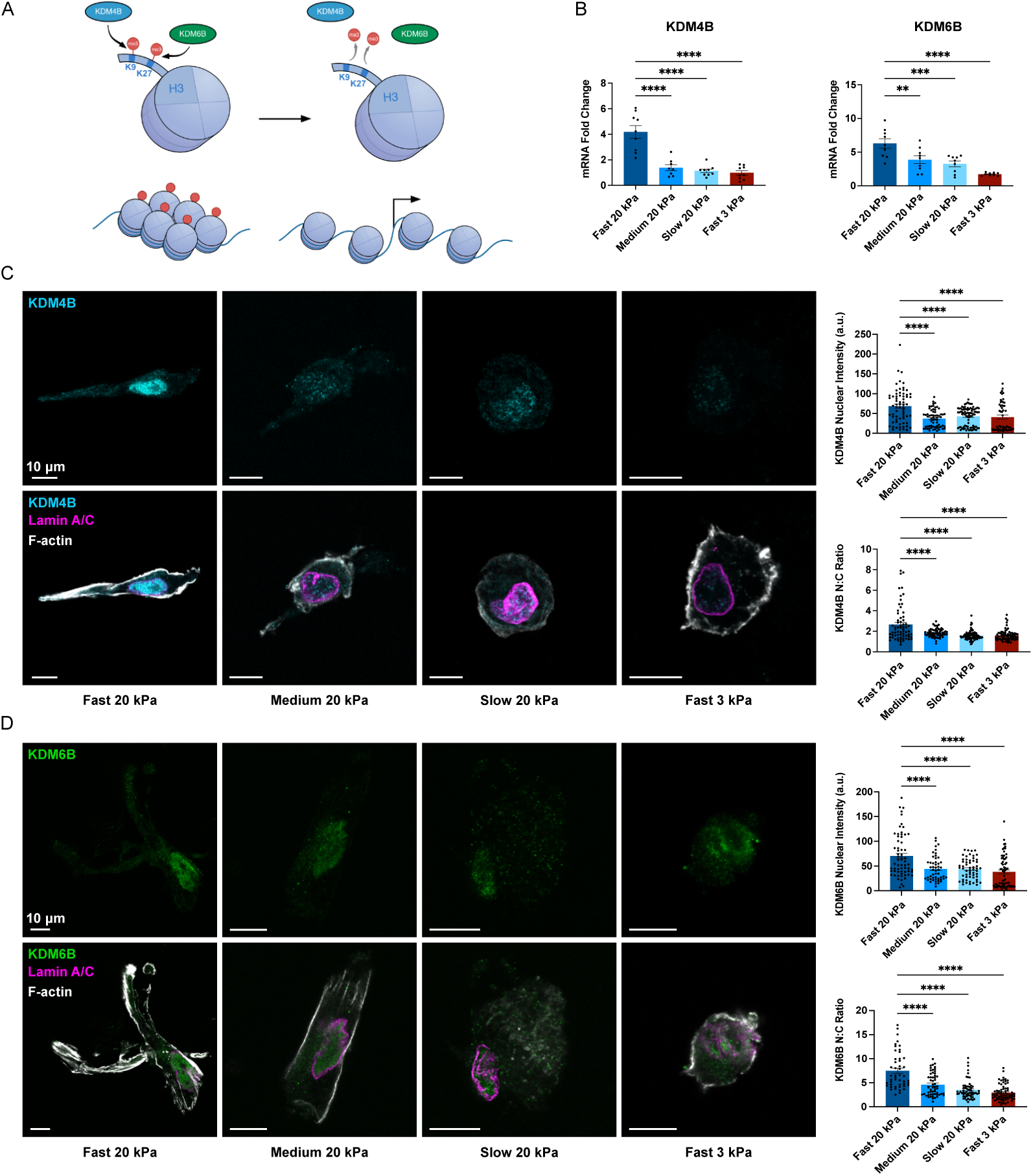
Stiff, fast-relaxing matrices promote expression of KDM4B and KDM6B. **A**, Schematic illustrating the role of KDM4B and KDM6B during chromatin remodeling. **B**, Gene expression of KDM4B and KDM6B between hydrogel conditions after seven days in culture. **C**, Immunofluorescence micrographs of KDM4B (left) and quantifications of KDM4B nuclear intensity and nuclear to cytoplasmic ratio (right). **D**, Immunofluorescence micrographs of KDM6B (left) and quantifications of KDM6B nuclear intensity and nuclear to cytoplasmic ratio (right). Statistical significance was determined by one-way analysis of variance (ANOVA) followed by Dunnett’s multiple testing correction. n = 3 replicates per condition. ** p < 0.01, *** p < 0.001, **** p < 0.0001.

H3K27 sites, respectively, we assessed the levels of these methylation post-translational modifications from cells across our four matrix conditions. Interestingly, we did not observe any significant differences in the total abundance of H3K9me3 or H3K27me3 after conducting western blot analysis (Supplemental Figure 2). However, western blotting can only capture global changes in methylation of these respective histone residues, which are potentially acted upon by other histone demethylases as well as methyltransferases. Our findings nonetheless show that matrix viscoelasticity promotes the expression and nuclear localization of KDM4B and KDM6B in an apparent dose-dependent manner.

### Inhibition of KDM4B and KDM6B reduces osteogenic differentiation in stiff, fast relaxing matrices

To interrogate the role of KDM4B and KDM6B during osteogenic lineage commitment, we encapsulated cells in stiff, fast-relaxing matrices, which induce the most robust osteogenesis, and used pharmacological antagonists to inhibit the catalytic domain of these demethylases. Treatment with inhibitors of KDM4B (ML-324, 10 µM) or KDM6B (GSK-J4, 15 µM) both resulted in significant decreases in osteogenic differentiation compared to the DMSO vehicle control, as assayed by quantification of ALP-positive cells (Figure 3A). Additionally, inhibition of KDM4B or KDM6B significantly decreased the expression of the early osteogenic markers RUNX2, ALP, and Collagen 1a1 (Figure 3B). Further, we observed a significant decrease in RUNX2 protein expression in both KDM4B and KDM6B-inhibited groups relative to the vehicle control (Figure 3C). Together, these results indicate that KDM4B and KDM6B are necessary for hBMSCs to undergo osteogenic differentiation in 3D viscoelastic matrices.

**Figure 3:**
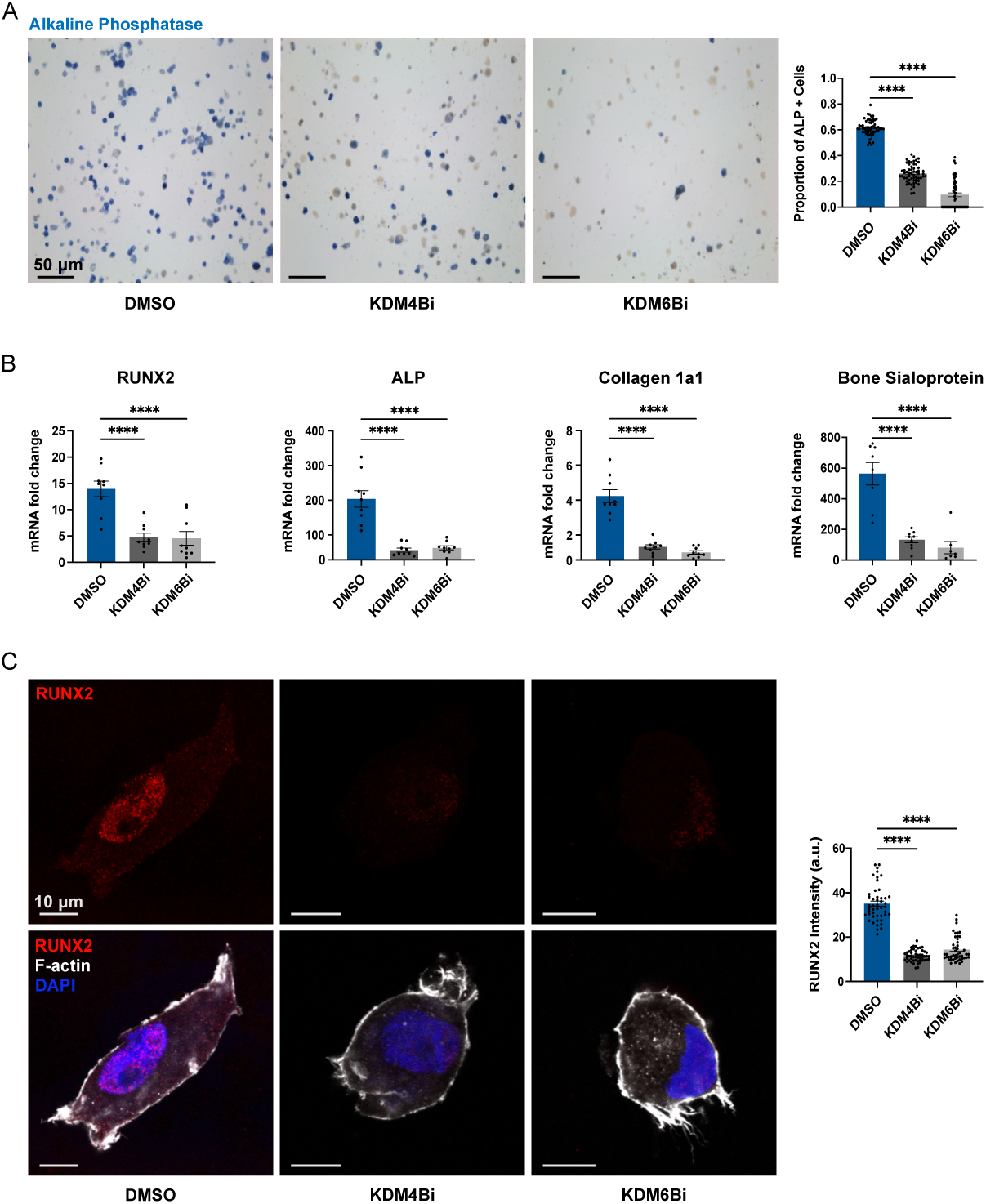
Inhibition of KDM4B or KDM6B reduces osteogenic differentiation in stiff, fast-relaxing matrices. **A,** Representative micrographs of stiff, fast-relaxing cryosections with alkaline phosphatase staining between DMSO-treated and small molecule-inhibited matrices (left), and quantifications of the proportion of ALP positive cells (right). **B**, Gene expression data for markers of early osteogenic differentiation between DMSO-treated and small molecule-inhibited gel groups. **C**, Representative immunofluorescence micrographs of cells stained for RUNX2 between DMSO-treated and small molecule-inhibited gel groups (left) and quantifications of RUNX2 staining intensity (right). Statistical significance was determined by one-way analysis of variance (ANOVA) followed by Dunnett’s multiple testing correction. n = 3 replicates per condition. **** indicates p < 0.0001.

### Inhibition of mechanotransduction pathway signaling reduces expression of KDM4B and KDM6B

Actin cytoskeletal contractility is a well-established mediator of MSC lineage-specific differentiation in response to environmental mechanical cues (Chen and Jacobs, 2013). In particular, Rho-associated kinase (ROCK)-mediated contractility has been shown to positively-regulate osteogenic differentiation of MSCs seeded on 2D substrates (McBeath *et al*., 2004) while exhibiting a concomitant inhibitory effect on adipogenic differentiation (Sordella *et al*., 2003). Similarly, non-muscle myosin II (NM2) activity has been shown to regulate osteogenic differentiation of MSCs in response to ECM stiffness (Engler *et al*., 2006). This contractility signaling occurs downstream of actin fiber bundling and focal adhesion kinase (FAK) activation, which are both influenced by integrin receptor clustering (Chaudhuri *et al*., 2016; Juhl IV *et al*., 2022). To assess the putative mechanotransduction pathways implicated in driving the expression of KDM4B and KDM6B, we perturbed cellular contractility by inhibiting ROCK (Y-27632, 10 µM), non-muscle myosin II (blebbistatin, 50 µM), or FAK (PF-573228, 10 µM). We observed significant reductions in the number of ALP-positive cells in response to treatment with each of these small molecule antagonists relative to the vehicle control, demonstrating the impact on hBMSC osteogenesis (Figure 4A). Inhibition of any of the three pathways led to a significant reduction in gene expression of Collagen 1a1, and inhibition of NM2 or FAK resulted in a significant decrease in gene expression of ALP (Figure 4B). Intriguingly, we found no significant differences in RUNX2 gene expression across these inhibitor treatments (Figure 4B). To determine whether KDM4B and KDM6B expression are regulated by these mechanosignaling pathways, we also assessed their expression by qPCR following inhibitor treatment. Expression of KDM4B was significantly decreased across all three mechanotransduction inhibitions (Figure 4B). Inhibition of ROCK or NM2 led to a significant decrease in the expression of KDM6B, though interestingly there was no significant difference in KDM6B expression upon FAK inhibition compared to the vehicle control. Importantly, these results illustrate that KDM4B and KDM6B expression is influenced by perturbations in mechanosignaling. To that end, this data indicates that early induction of osteogenic gene expression is not necessarily predicated upon the same mechanically actuated pathways, and instead suggests that MSCs utilize numerous, orthogonal force transduction pathways during osteogenic lineage commitment.

**Figure 4:**
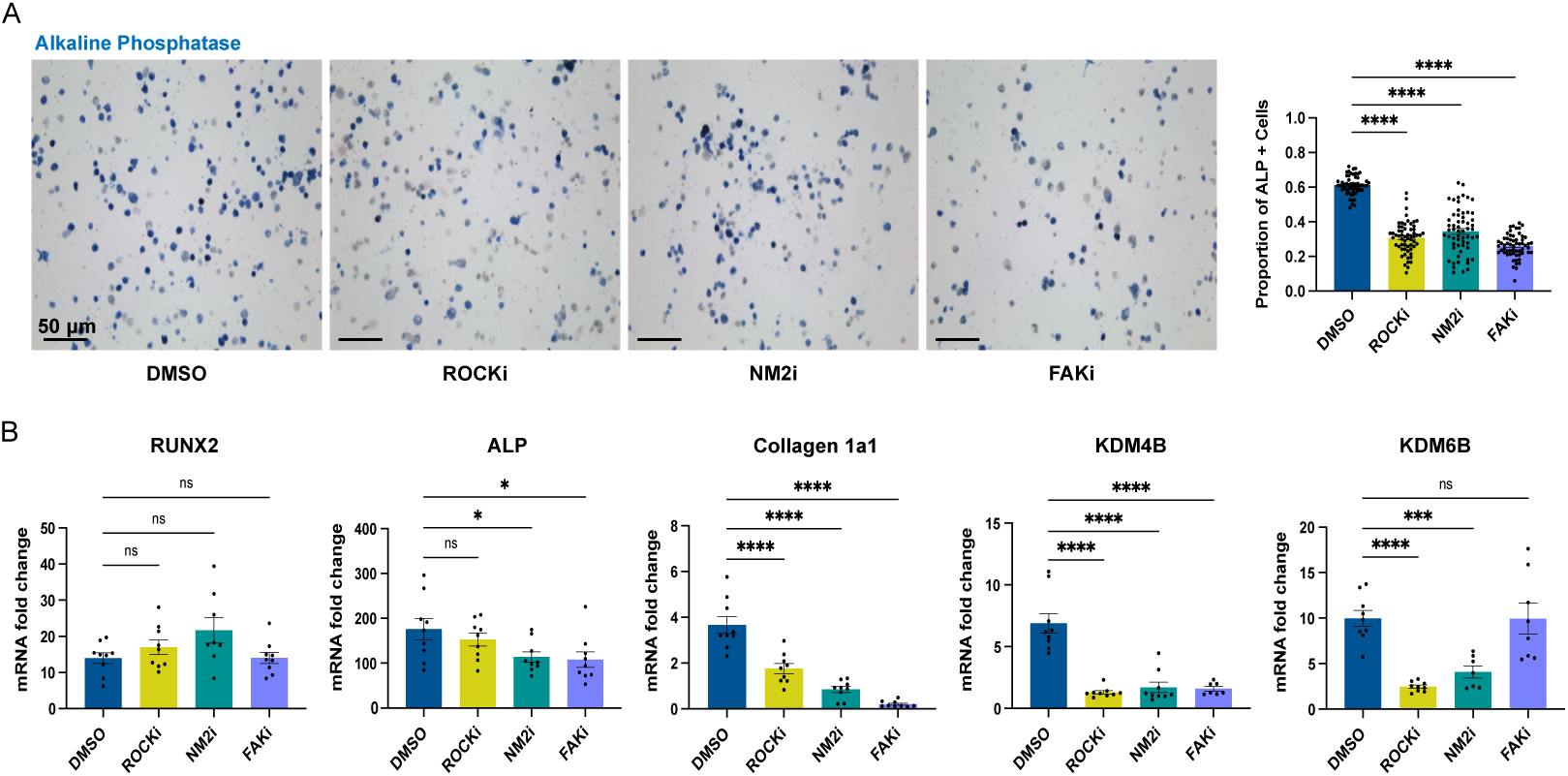
Inhibition of mechanotransduction pathways reduces expression of KDM4B and KDM6B. **A,** Representative micrographs of ALP stained stiff, fast-relaxing matrices comparing DMSO-treated and small molecule-inhibited conditions(left), and quantifications of ALP positive cells (right). **B**, Gene expression data for osteogenic markers as well as KDM4B and KDM6B comparingDMSO-treated and small molecule-inhibited conditions. Statistical significance was determined by one-way analysis of variance (ANOVA) followed by Dunnett’s multiple testing correction. n = 3 replicates per condition. * indicates p < 0.05, ** p < 0.01, *** p < 0.001, **** p < 0.0001.

### Stiff, fast-relaxing matrices increase SMAD 1/5/8 activation and KDM expression

BMP signaling governs osteogenic differentiation through activation of BMP Type 1 and 2 receptors (Urist, 1997). The cytosolic kinase domains of these receptors induce reciprocal phosphorylation of one another, which then activates these domains to phosphorylate downstream effectors such as SMADs 1/5/8. Upon phosphorylation of SMAD 1/5/8 (pSMAD 1/5/8), these transcription factors translocate to the nucleus and promote expression of osteogenic-related genes (Beederman *et al*., 2013; Rahman *et al*., 2015; Wu *et al*., 2016). Prior research has shown that BMP 4/7 treatment significantly increases KDM4B and KDM6B expression in BMSCs, and that SMAD 1 knockdowns in these same cells led to a decrease in KDM4B and KDM6B expression (Ye *et al*., 2012). Accordingly, we were curious to investigate the impact of stiffness and stress relaxation on SMAD 1/5/8 activation. We observed the highest abundance of pSMAD 1/5/8 in cells cultured in stiff, fast-relaxing gels via immunofluorescence imaging (Figure 5A). The differences in pSMAD 1/5/8 abundance were significantly greater in stiff and fast-relaxing matrices compared to slower relaxing or soft matrices. To investigate the contribution of SMAD 1/5/8 signaling during osteogenic differentiation in stiff, fast-relaxing matrices, we used LDN-193189 (0.1 µM), a small molecule that specifically blocks BMP Type 1 Receptor (BMPR1) phosphorylation of SMAD 1/5/8 (Boergermann *et al*., 2010). Upon treatment of hBMSCs in stiff, fast-relaxing matrices with LDN-193189 for seven days, we saw a significant decrease in ALP-positive cells relative to the vehicle control (Figure 5B). Inhibition of BMPR1 also led to a significant decrease in gene expression of osteogenic markers RUNX2 and ALP (Figure 5C). Importantly, inhibition of BMPR1 phosphorylation of SMAD 1/5/8 also resulted in significant decreases in gene expression of KDM4B and KDM6B, respectively (Figure 5C). Furthermore, this inhibition also led to a significant decrease in both the nuclear localization and abundance of KDM4B and KDM6B relative to the vehicle control (Figure 5D-E). These results illustrate a linkage between BMP signaling and the expression of KDM4B and KDM6B, providing insight into how matrix mechanical cues influence BMP receptor activity during osteogenic lineage commitment.

**Figure 5:**
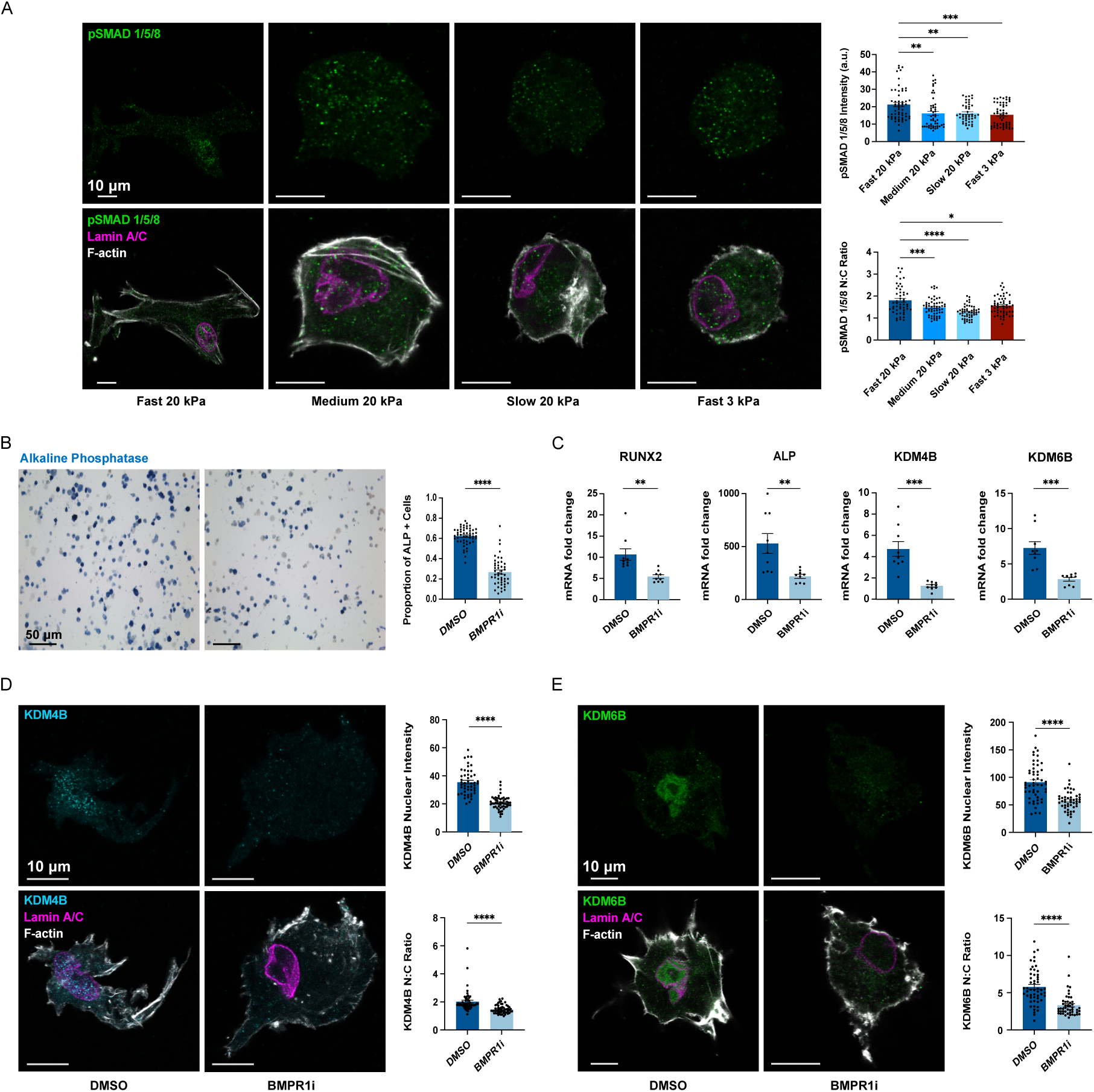
Viscoelasticity increases BMP-SMAD pathway signaling and regulates expression of KDM4B and KDM6B. **A**, Representative micrographs of cells cultured across matrix conditions stained for phospho-SMAD 1/5/8, lamin A/C, and F-actin (left) with quantifications of signal intensity and nuclear to cytoplasmic ratio (right). **B**, Alkaline phosphatase staining of cells in stiff,fast-relaxing matrices comparing DMSO-treated and BMPR1-inhibited media conditions (left) with quantification of ALP positive cell proportions. **C**, Gene expression quantifications of DMSO-treated and BMPR1-inhibited conditions. **D-E**, Representative micrographs of cells stained for KDM4B, KDM6B, lamin A/C, and F-actin between DMSO-treated and BMPR1-inhibited conditions with quantifications of nuclear to cytoplasmic ratio and nuclear intensity. For panel A, statistical significance was determined by one-way analysis of variance (ANOVA) followed by Dunnett’s multiple testing correction. For panels B-E, statistical significance was determined by unpaired t tests with Welch’s correction. n = 3 replicates per condition. * indicates p < 0.05, ** p < 0.01, *** p < 0.001, **** p < 0.0001.

## Discussion

In this study, we investigated the mechanobiological regulation of histone demethylases KDM4B and KDM6B during early osteogenic differentiation in human BMSCs. We observed that matrix stiffness and stress relaxation rate modulate the expression and nuclear localization of both KDM4B and KDM6B, and that inhibition of several mechanotransduction pathways decreases the expression of these demethylases. Our findings represent the first investigation of how SMAD 1/5/8 activation is influenced by matrix viscoelasticity and provides new insight into how BMSCs integrate ECM mechanical cues to coordinate lineage specification. To that end, our results also contribute to an emerging field investigating how matrix mechanics shape cell fate outcomes through epigenetic regulation by histone modifying enzymes.

Recent findings in the field of mechanobiology have illustrated an interplay between mechanotransduction signaling and epigenetic remodeling during phenotypic transitions across a variety of mesenchymal-derived cell types in direct response to microenvironmental cues (Downing *et al*., 2013; Heo *et al*., 2015, 2023; Roy *et al*., 2018, 2020; Wu *et al*., 2025). In the context of disease onset, such as gastric and mammary tumorigenicity, changes in ECM mechanics have been linked to aberrations in gene expression specifically through changes in DNA methylation, chromatin accessibility and HDAC3 and HDAC8 activity (Stowers *et al*., 2019; Jang *et al*., 2021).

Interestingly, hallmarks of osteogenic differentiation in BMSCs have been shown to correlate with both changes in expression of histone deacetylases and associated histone acetylation in response to stiff substrates (Killaars *et al*., 2019, 2020). However, the impact of stress relaxation on epigenomic changes during stem cell differentiation has not been studied to date. Our findings contribute to the burgeoning effort to understand how ECM mechanical properties both drive changes in cell fate through the activity of epigenetic signaling pathways.

Our results corroborate previous reports that KDM4B and KDM6B are essential for the induction of osteogenesis and that loss of KDM4B and KDM6B significantly reduces expression of early osteogenic markers (Ye *et al*., 2012; Deng *et al*., 2022; Jin *et al*., 2022; Ying *et al*., 2024). In line with our observations that inhibiting the catalytic activity of these demethylases reduces expression of essential osteogenic markers, previous findings have identified that KDM4B removes repressive methylation marks from the promoter of RUNX2 to facilitate its transcription and establish osteogenic lineage commitment (Kang *et al*., 2022), underscoring the necessity of these demethylases to initiate osteogenic differentiation. RUNX2 is considered a master regulator of osteogenic differentiation, and this evidence relating KDM4B demethylation activity to the expression of RUNX2 suggests a molecular mechanism through which KDM4B is essential for early osteogenesis. Similarly, KDM6B depletion has been shown to significantly reduce ALP activity and associated mineral deposition (Xu *et al*., 2013). Thus, our work adds to the literature showing numerous, orthogonal signaling pathways that promote osteogenesis, including via epigenomic remodeling by histone modifiers such as KDM4B and KDM6B.

Matrix mechanics are known to regulate osteogenesis, and significant work in the field of mechanobiology has demonstrated that mechanotransduction pathways are essential for osteogenic differentiation. Our use of pharmacological inhibitors targeting downstream effectors of integrin signaling, including ROCK, NM2, and FAK, provide novel evidence that KDM4B and KDM6B are sensitive to various force transduction modalities. While expression of both KDM4B and KDM6B were significantly decreased upon inhibiting ROCK and NM2, RUNX2 expression did not change significantly in response to these inhibitors. MSCs have long been understood to differentiate based on actomyosin-driven signaling (Engler *et al*., 2006), but prior work has shown that RUNX2 mRNA levels do not change upon ROCK inhibition (Prowse *et al*., 2013) and that numerous upstream regulatory processes may govern its expression beyond cytoskeletal contractility (Franceschi *et al*., 2003). Reports have also shown that FAK inhibition significantly reduces both ALP and Collagen 1a1 expression in BMSCs cultured in 2D (Rajshankar *et al*., 2017; Gunn *et al*., 2022), which corroborates with our observations of this inhibition in stiff, fast-relaxing 3D matrices. While we observed a significant decrease in KDM4B expression upon inhibiting FAK, this inhibition did not lead to a significant difference in KDM6B mRNA levels. Whether BMSCs utilize orthogonal pathways to activate KDM4B and KDM6B during mechanically induced osteogenesis is an open question with promise for future investigation.

Smad 1/5/8 activation is known to regulate osteogenesis of BMSCs (Wu *et al*., 2016). Recent findings have suggested that stiff matrices augment osteogenesis by potentiating activity of the BMP-SMAD 1/5/8 signaling axis via focal adhesion signaling (Görlitz *et al*., 2024). Additionally, inhibition of actomyosin contractility has been shown to decrease the response of BMSCs to BMP signaling in 3D culture. (Rath *et al*., 2011; Li *et al*., 2024b). Our findings indicate that both stiffness and stress relaxation exert an effect on SMAD 1/5/8 signaling, and that this pathway regulates the expression of histone modifiers KDM4B and KDM6B, as well as canonical osteogenic markers RUNX2 and ALP. A recent report has also shown that collagen alignment upregulates SMAD 1/5/8 signaling in MSCs in the absence of BMP ligands onto cells (Heo *et al*., 2025), where the alignment of collagen fibers within the ECM promotes cell spreading and mechanosignaling.

Despite the lack of collagen or any fibrillar ECM component, we also found similar significant increases in cell spreading and the degree of osteogenesis in stiff, viscoelastic matrices.

Additionally, another report showed that disruptors of either actin polymerization or myosin ATPase activity negatively regulates SMAD 1/5/8 activation and precludes robust osteogenic differentiation (Xu *et al*., 2017). This evidence highlights a potential overlap in canonical BMP signaling with biophysical elements of the cellular microenvironment, and that BMP-regulation of osteogenesis is predicated, in part, on cell-ECM interactions. This convergence of both biochemical and mechanical regulation of SMAD 1/5/8 activity may therefore serve as the target of future investigations toward designing biomaterial platforms to study bone regeneration in the context of critical wound healing and age-related bone loss.

Our work shows a novel mechanoresponsive role for KDM4B and KDM6B in regulating BMSC osteogenesis. Specifically, we found that the expression and nuclear abundance of KDM4B and KDM6B is enhanced in stiff and fast-relaxing matrices during osteogenesis. These results illustrate how both matrix stiffness and stress relaxation regulate BMSC fate via activation of mechanosignaling pathways and further, that the SMAD 1/5/8 pathway is upregulated in stiff, viscoelastic matrices. In conclusion, this work aims to better understand how ECM mechanics are linked to epigenetic regulation and how further delineation of these signaling pathways can be leveraged to guide stem cell fate for applications in both disease modeling and regenerative medicine.

## Materials and Methods

### Hydrogel Preparation

Pronova VLVG and LF 20/40 alginate were purchased from NovaMatrix and prepared as previously described (Rowley *et al*., 1999; Chaudhuri *et al*., 2016). Briefly, RGD–modified alginate was prepared by conjugating the oligopeptide GGGGRGDSP (Peptide 2.0) to the alginate polymers using carbodiimide chemistry at a final concentration of 1,500 μM RGD in 2% wt/vol alginate hydrogels. Alginate was then dialyzed against deionized water for 2–3 days (molecular weight cutoff of 10 kDa), treated with activated charcoal, sterile filtered, lyophilized, and then reconstituted in serum-free Dulbecco’s Modified Eagle Medium (DMEM, Life Technologies).

Polyethylene glycol (PEG)–alginate was prepared by coupling PEG-amine (5 kDa, Laysan Bio) to VLVG alginate using carbodiimide chemistry and purified with a similar procedure to the RGD coupling.

### Mechanical Characterization

Rheology measurements were conducted using a stress-controlled Anton Paar MCR 502 rheometer. Hydrogels were cast between silanized glass plates with a 2 mm spacer, biopsy punched into 8 mm diameter discs and allowed to swell in DMEM overnight. The storage modulus (G’) and loss modulus (G’’) of alginate gels were measured using at a frequency of 1 Hz with 0.1% applied shear strain. The complex modulus (G*) was measured and used to calculate the elastic modulus (E), as shown below. The Poisson’s ratio (*v*) was assumed to be 0.5 (Charbonier *et al*., 2021).

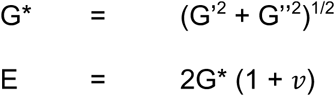

Stress relaxation tests were conducted using a constant strain of 15%. The time required for each gel sample to reach half of the maximum stress was used to determine the stress relaxation time (τ_1/2_) for each gel condition across three independent replicates. Gel recipes are detailed in Table S1.

### Cell Culture

Primary human BMSCs (hBMSCs) were obtained from Millipore Sigma (SCC034) and cultured according to established protocols (Huebsch *et al*., 2010; Darnell *et al*., 2017). Cells were expanded in low-glucose DMEM in the presence of 20% FBS (ThermoFisher) and 1% Pen/Strep (Gibco) and maintained below 80% confluency.

### Cell Encapsulation and Osteogenic Differentiation

hBMSCs used in this study were expanded up to passage 4 prior to encapsulation in hydrogel matrices. Cells were washed twice with Dulbecco’s phosphate-buffered saline (DPBS) prior to trypsinization in 0.25% trypsin/EDTA (Gibco). Cells were then suspended in DMEM and mixed with alginate. The alginate-cell solution was mixed with CaSO_4_ and immediately cast between two glass plates. Upon gelation, gels were punched using 8 mm biopsy punches and cultured in OsteoMAX differentiation media (Millipore Sigma, SCM121) for 7 days with media changes occurring every 2 days.

### Immunofluorescence Staining

Encapsulated cells were fixed in 4% paraformaldehyde (PFA) for 45 min and then washed three times in DPBS with Ca^2+^/Mg^2+^ for 15 min each. The hydrogels were then dehydrated in a 30% sucrose solution in DPBS overnight and then incubated in a 1:1 mixture of 30% sucrose and OCT solution (ThermoFisher, 23730571) prior to freezing on dry ice. Frozen hydrogel samples were cryosectioned into 40 µm-thick slices and were adhered to poly lysine-coated slides for subsequent staining procedures. Slides were blocked with a solution of 10% goat serum (ThermoFisher), 1% bovine serum albumin (Sigma), 0.1% Triton X-100 (Sigma) and 0.3 M glycine (Sigma) for 1 h at room temperature. Primary antibodies were diluted in the blocking solution and incubated at 4°C overnight. The slides were then washed three times in DPBS for 5 min each prior to being incubated with Alexa Fluor 488, 555, or 647 secondary antibodies in blocking solution for 1 h at room temperature. Alexa Fluor 488 or 647–phalloidin (1:100 dilution, ThermoFisher) and 4′,6-diamidino-2-phenylindole (DAPI, 1 µg ml^−1^, ThermoFisher) were also diluted in the blocking solution and simultaneously incubated with secondary antibodies for 1 h. The slides were then washed three times for 5 min in DPBS, and coverslips were applied with Prolong Gold antifade reagent (ThermoFisher, P36930). Antibodies used in this study include Integrin β1 (ThermoFisher, 14-0299-82), Paxillin (Cell Signaling Technologies, 50195S), KDM4B (Cell Signaling Technologies, 8639), KDM6B (Abcam, ab169197), RUNX2 (Cell Signaling Technologies, 12556S), Lamin A/C (Cell Signaling Technologies, 4777S), and phospho-SMAD 1/5/8 (Cell Signaling Technologies, 13820). Slides were imaged on a Leica SP8 laser scanning confocal microscope with a 63x objective.

### ALP staining

ALP was stained using Fast Blue BB Salt (Millipore Sigma, F3378) and naphthol-AS-MX phosphate (Millipore Sigma, N4875) at a working concentration of 500 µg/ml each in an alkaline buffer (100 mM Tris-HCl, 100 mM NaCl, 0.1% Tween-20, 50 mM MgCl2, pH = 8.2). Poly lysine-coated slides with affixed cryosections were washed three times in DPBS, equilibrated in alkaline buffer for 15 min, then incubated in FastBlue working solution for 60 min at room temperature. The slides were then washed for 15 min in alkaline buffer followed by three washes in DPBS for 5 min each and mounted using Prolong Gold antifade reagent. Slides were imaged using an Olympus BX60 Upright microscope using a color camera and 10x objective. The number of ALP-positive cells was enumerated and then divided by the total the number of cells in each field of view to determine the total proportion of ALP-positive cells.

### Image Analysis

FIJI (NIH) was used for image analysis to quantify cell area, circularity, localization, and intensity of the selected markers used in this study. The plugin Ilastik was used to analyze confocal microscopy images and generate subcellular region-specific ROIs that assign labels to pixels based on their unique features and user annotations. The generated ROIs were imported into ImageJ, where a custom macro was used to threshold, analyze, and create a mask of defined cell regions. The quality of the mask was manually verified before defining a Region of Interest (ROI) and measuring the fluorescent intensity.

To determine the nuclear-to-cytoplasmic signal ratio, masks were generated for the cell nucleus (lamin A/C) and cell outline (phalloidin). The nuclear mask was subtracted from the cell outline mask to generate a ROI for the cytoplasm, then both the cytoplasm and nuclear ROI were measured for the protein channel of interest. The mean fluorescence intensity in each ROI was calculated by dividing the total fluorescence intensity by the number of pixels measured.

Focal adhesions were identified by the colocalization of paxillin and integrin β1. To assess this, the same Ilastik pipeline was used to generate cell outline masks based on F-actin staining. The ImageJ plugin *ComDet* was then used to quantify the degree of colocalization within each cell ROI, using an average particle size of 8 pixels, a maximum distance of 6 pixels between the centers of colocalized spots, and an intensity threshold set to 20 standard deviations above the mean for both paxillin and integrin β1 channels. Additionally, the mean fluorescence intensity of paxillin was measured within each cell ROI.

### Quantitative real-time PCR

Cells were extracted from hydrogels by chelation of calcium ions using 50 mM EDTA in DPBS. Cell pellets were washed twice with DPBS to remove residual EDTA. Cells were then lysed with Trizol (Life Technologies, 10296028) and RNA was extracted according to the manufacturer’s instructions (Epoch Life Sciences, 1660250). Purified RNA was then reverse-transcribed into complementary DNA (cDNA) using a High-Capacity cDNA Reverse Transcription Kit (Applied Biosystems™, 4368814). Here, 10 ng of cDNA was used for each quantitative real-time PCR (qRT-PCR) reaction, performed in triplicates for each biological replicate. SYBR Green PCR Master Mix (ThermoFisher, A25742) was used for amplification of target sequences on a CFX96 Touch Real Time Detection System (Bio-Rad). Relative gene expression was calculated using the 2^−ΔΔCt^ method (Livak and Schmittgen, 2001). The primer sequences are detailed in Table S2.

### Western blotting

Cells were extracted from hydrogels by chelation of calcium ions using 50 mM EDTA in DPBS. Once pelleted, cells were washed twice with DPBS to remove any residual EDTA. Cells were then lysed via sonication for 30 seconds in RIPA buffer (ThermoFisher, 89901) along with protease & phosphatase inhibitor cocktail (ThermoFisher, 78440). Protein lysates were centrifuged to pellet cell debris, and the supernatant was collected and used for further analysis. Protein samples were separated using SDS-PAGE and transferred to nitrocellulose membranes. Membranes were blocked in Tris-buffered saline (TBS) with 5% w/v nonfat milk and incubated with primary antibodies for H3K9me3 (Cell Signaling, D4W1U), H3K27me3 (Cell Signaling, C36B11), and H3 (ThermoFisher, 39763) at 4°C overnight. Membranes were then washed with TBS with 0.05% Tween-20 and incubated with Alexa Fluor-conjugated IgG secondary antibodies (ThermoFisher) for 1 h. Protein bands were imaged on a ChemiDoc system (Bio-Rad).

### Pharmacological Inhibition

For inhibition experiments, small molecule antagonists of KDM4B (ML-324, 10 µM) and KDM6B (GSK-J4, 15 µM) were added to the differentiation media for culture over 7 days. For inhibition of mechanotransduction signaling, small molecule antagonists of ROCK (Y-27632, 10 µM), non-muscle myosin II (Blebbistatin, 50 µM), and FAK (PF-573228, 10 µM) were added to the differentiation media for culture over 7 days. To inhibit BMP Type 1 Receptor (BMPR1), the small molecule antagonist, LDN-193189 (0.1 µM), was added to the differentiation media for culture over 7 days. All inhibitors were purchased from Cayman Chemical and dissolved in dimethyl sulfoxide (DMSO).

### Statistical analysis

All statistical analyses were conducted using GraphPad Prism 10.4. All statistical tests performed for analysis are described in each respective figure caption.

## Acknowledgements

We thank members of the Stowers lab for critical feedback on this manuscript. We acknowledge the use of the NRI-MCDB Microscopy Facility at University of California, Santa Barbara. Furthermore, we acknowledge the use of the Biological Nanostructures Laboratory within the California NanoSystems Institute, supported by the University of California, Santa Barbara and the University of California, Office of the President. A.Z. was supported by the California Institute of Regenerative Medicine (CIRM) COMPASS EDUC5-13744 training grant.

## Author contributions

Conceptualization, I.M.T., R.S.S.; methodology, I.M.T., R.S.S.; validation, I.M.T., A.Z.; formal analysis, I.M.T., A.Z.; investigation, I.M.T., A.Z.; resources, A.Z., R.S.S.; writing – original draft, I.M.T., R.S.S.; writing – review and editing, I.M.T., A.Z., R.S.S.; visualization, I.M.T., A.Z., R.S.S.; supervision, R.S.S.; funding acquisition, R.S.S.

