## Supplemental Materials for "Matrix stiffness and stress relaxation regulate osteogenesis through histone demethylases KDM4B and KDM6B"

**Supplemental Table 1:** Alginate hydrogel recipes for all conditions included in this study.

| **Matrix Condition** | **Sodium Alginate** | **Alginate MW** | **$Ca^{2+}$ conc.** |
|---|---|---|---|
| Fast 20 kPa | 2% UPVLVG-PEG | < 75 kDa | 80 mM |
| Medium 20 kPa | 2% UPVLVG | < 75 kDa | 55 mM |
| Slow 20 kPa | 2% LF20/40 | > 200 kDa | 18 mM |
| Fast 3 kPa | 2% UPVLVG-PEG | < 75 kDa | 32 mM |

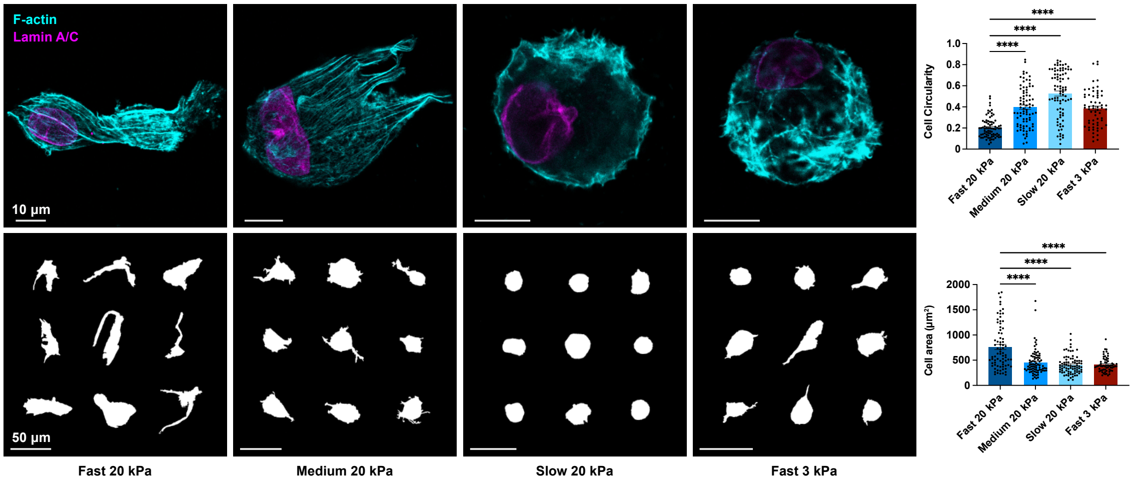


**Supplemental Figure 1: Cell spreading increases with stiffness and stress relaxation in 3D.** Representative micrographs of cells cultured across hydrogel conditions with associated representative cell outlines (left), and quantifications of both cell circularity and cell area (right). Statistical significance was determined by one-way analysis of variance (ANOVA) followed by Dunnett's multiple testing correction. **** $p < 0.0001$.

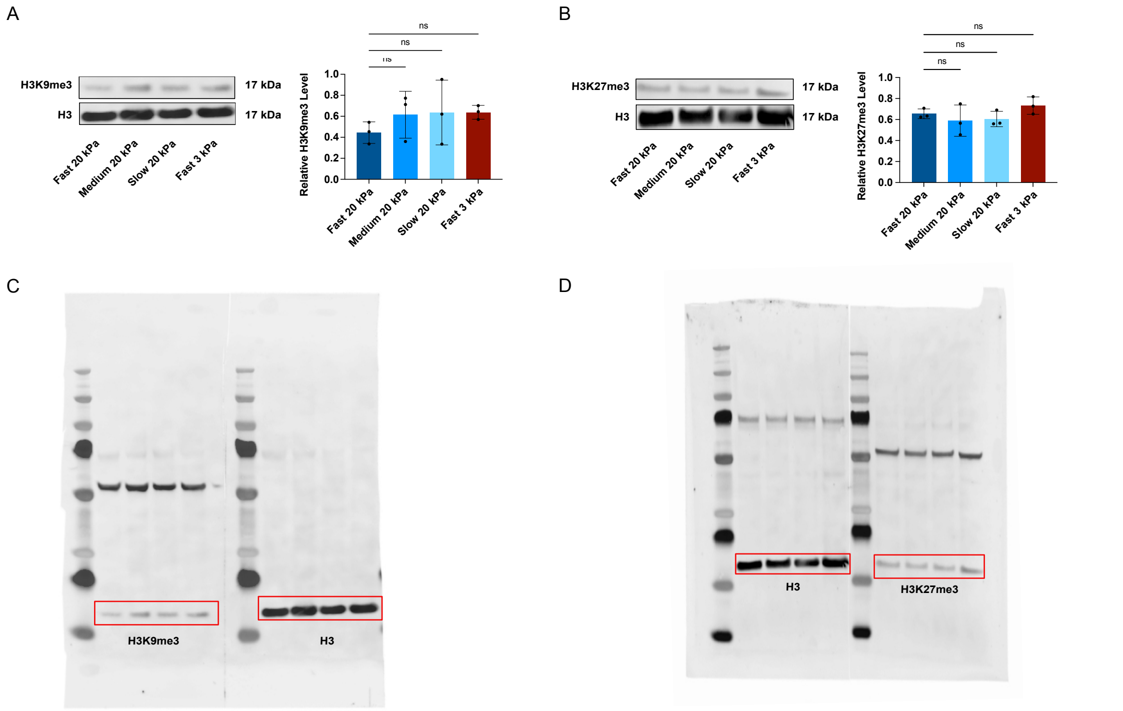


**Supplemental Figure 2: A-B**, Representative western blots for H3K9me3, H3K27me3, and loading control H3 for hBMSCs cultured across hydrogel conditions for seven days with associated quantifications. **C-D**, Whole blots from which the lanes displayed in panels A-B were cropped. Statistical significance was determined by one-way analysis of variance (ANOVA) followed by Dunnett's multiple testing correction. ns where $p > 0.05$.

**Supplemental Table 2:** Primer sequences used for qPCR experiments.

| Gene target | Forward Primer | Reverse Primer |
|---|---|---|
| RUNX2 | GACCAGTCTTACCCCTCCTAC | CTGCCTGGCTCTTCTTACTGA |
| ALP | ACTCCCACTTCATCTGGAAC | CCTGTTCAGCTCGTACTGCA |
| Collagen 1a1 | GAGAGGAAGGAAAGCGAGGAG | GGGACCAGCAACACCATCT |
| Osteocalcin | TGACGAGTTGGCTGACCA | AGGGTGCCTGGAGAGGAG |
| Bone Sialoprotein | TGCCTTGAGCCTGCTTC | GCAAAATTAAAGCAGTCTTCATT |
| KDM4B | GGACTAGAGGCCGTCTAAATTG | ACTTCCTGCGTGCAAAGA |
| KDM6B | CACGCGGCTCGTGTATGTA | GGGTCACAGCTAGCATTGGAA |
| GAPDH | ACAGCGACACCCACTCCT | GAGGTCCACCACCCTGTT |
